# The importance of M1 muscarinic receptor phosphorylation in learning and memory

**DOI:** 10.64898/2026.03.23.713145

**Authors:** Aisling McFall, Colin Molloy, Katharine Gibson, Craig W. Lindsley, Andrew B. Tobin

**Affiliations:** Centre for translational pharmacology, School of Molecular Biosciences, University of Glasgow; School of Cardiovascular and Metabolic Health, University of Glasgow; Department of Pharmacology, Vanderbilt University

## Abstract

The muscarinic acetylcholine receptor 1 (mAChR1, M1) has been identified as a primary target for Alzheimer’s disease (AD) and better understanding of the receptor biology, especially in regard to biased signalling of the receptor, will allow for the development of improved drugs targeting cholinergic dysfunction in AD. The aim of this study was to determine the contribution of phosphorylation of M1 to the learning and memory (LM) effects of M1 agonism.

The contribution of M1 phosphorylation dependent signalling in LM was assessed using the mAChR1 positive allosteric modulator, VU0486846, in a scopolamine (1.5 mg/kg) induced LM deficit model in mice expressing HA-tagged M1 (M1-WT), phosphorylation deficient HA-tagged M1 (M1-PD), or mice deficient in M1 (M1-KO). LM was assessed using a fear conditioning (FC) testing paradigm where context and cued memory retrieval was measured 24 hrs after training and a higher level of freezing indicated intact memory. Results demonstrated that scopolamine induced a significant LM deficit in both context and cued retrieval in M1-WT mice which was partially rescued by VU0486846 confirming a contribution of M1 signalling in LM. The scopolamine induced deficit in contextual retrieval in M1-KO mice was not rescued by VU0486846, which is an M1 selective ligand, while scopolamine did not induce a deficit in cued retrieval in M1-KO mice. In M1-PD mice, scopolamine induced a LM deficit in contextual retrieval, however this was also not rescued by VU0486846 treatment. Similarly to M1-KO animals, M1-PD mice did not display a scopolamine induced deficit in cued retrieval. When freezing responses were compared across strains, M1-PD mice displayed a deficit compared to M1-WT and M1-KO mice in contextual retrieval, while both M1-PD and M1-KO mice displayed a deficit compared to M1-WT mice in cued retrieval. These results demonstrate that although M1 agonism can restore a LM deficit in both contextual and cued testing paradigms, only the cued retrieval response is dependent on the M1. Additionally, biased G_q_ M1 signalling is not sufficient to restore contextual memory and requires phosphorylation of the receptor. Furthermore, biased M1 signalling results in LM deficits not seen with KO of the receptor. Overall, these results reiterate the importance of considering the bias of ligands when developing M1 agonists for dementia in the future.

## Introduction

The cholinergic hypothesis of Alzheimer’s disease (AD) poses that a lack of cholinergic transmission is one cause of the cognitive deficits displayed in the disease. As a result, acetylcholinesterase inhibitors, such as donepezil, to increase levels of endogenous acetylcholine (ACh), are commonly prescribed for symptomatic management of AD. Unfortunately, however, due to systemic increases in levels of ACh, these drugs are associated with adverse dose-limiting side effects and therefore their efficacy is limited. ACh is a neurotransmitter which acts on either nicotinic acetylcholine receptors (nAChR) or muscarinic acetylcholine receptors (mAChR). One alternative therapeutic avenue is to directly target the mAChR1 (M1 receptor) as this is the most highly expressed mAChR in the brain; indeed up to 60% of the muscarinic receptors within the human and mouse brain are this subtype (Bradley et al., 2017; Flynn et al., 1995). In preclinical studies, targeting of M1 has demonstrated reduced amyloid beta levels (Beach et al., 2001), enhanced cognition (Ragozzino et al., 2012; Rook et al., 2018) and disease modifying effects in models of neurodegeneration (Abd-Elrahman et al., 2022; Bradley et al., 2017). Meanwhile, in clinical trials, the M1/M4 preferring agonist xanomeline, failed due to adverse effects, however this was attributed to off-target effects at the M2 and M3 receptors. Subsequently, a structure-based drug design approach yielded a specific M1/M4 agonist, HTL9936, which showed a promising safety profile and activity in the cognitive centres of human volunteers (Brown et al., 2021). To circumnavigate the difficulties in targeting the highly conserved orthosteric site, positive allosteric modulators (PAMs) provide an alternative option for specific targeting of the M1 mAChR. PAMs can either be “pure” PAMs, meaning they enhance the pharmacological effects of bound endogenous orthosteric ligand, or PAM-agonists, which have intrinsic agonist activity independent of the enhancement effect. Several M1-PAMs have been developed and have demonstrated rescue of cognitive deficits in rodent models, however many of these are actually PAM-agonists and can induce adverse effects due to off-target M1 activation rather than endogenous ACh signalling enhancement (Dwomoh, Tejeda, et al., 2022). VU0486846 (VU846) is a promising “pure” PAM which has showed efficacy in cognition (Rook et al., 2018) and disease modifying effects in rodent models of neurodegeneration (Abd-Elrahman et al., 2022; Bradley et al., 2017; Dwomoh, Rossi, et al., 2022). Vanderbilt University have subsequently developed a more advanced PAM, VU319, which has showed promising results in terms of safety and cognitive performance in human volunteers (Conley et al., 2021; Newhouse et al., 2020).

Evidently, targeting the M1 mAChR shows promise for the treatment of AD and improvement in cognition; however, a better understanding of the biology of this receptor is important for successful translation of therapies. The mAChR family are G-protein coupled receptors (GPCRs), with M1 specifically being coupled to G_q/11_. Like many other GPCRs, M1 undergoes rapid phosphorylation, desensitisation and internalisation upon agonist stimulation (Waugh et al., 1999) and, indeed, phosphorylation of M1 may be utilised as a marker of receptor activation following learning and memory (LM) events (Butcher et al., 2016). While we have previously demonstrated that activation of M1, with an M1-PAM, improves LM and survival in a murine prion model of terminal neurodegeneration (Bradley et al., 2017), we have also shown that phosphorylation of M1 is important for neuroprotection with mice expressing a phosphorylation-deficient M1 receptor (M1-PD) displaying worsened disease pathology than M1-WT mice (Scarpa et al., 2021). In addition, we have demonstrated that lack of phosphorylation of M1 results in a high anxiety phenotype and may impair working memory (Bradley et al., 2020), however a detailed study assessing the importance of phosphorylation of the M1 mAChR in LM is lacking. Therefore, here we aimed to utilise M1-PD mice to establish if phosphorylation of M1 is required for the LM effects of M1 agonism and assess how M1-PD mice compare to both M1-WT and M1-knockout (M1-KO) animals using context and cue dependent fear memory testing.

## Results

### Drug dose optimisation

Scopolamine induced an LM deficit in terms of contextual memory retrieval when administered to M1-WT mice at doses from 1 – 3 mg/kg 30 mins prior to fear conditioning. This deficit was most robust at a dose of 1.5 mg/kg scopolamine with a reduction in context dependent freezing from 53.8% ± 8.3% to 34.0% ± 14.7% (Figure 1A). In a separate experiment, this LM deficit induced by 1.5 mg/kg scopolamine (freezing level vehicle: 57.5%± 23.4%, scopolamine: 37.5% ±19.5%) was rescued by coadministration of the M1-PAM VU846 at a dose of 10 mg/kg (freezing level 52.1% ±12.3%) but not the higher doses of 30 or 100 mg/kg (freezing level 36.1% ± 17.4% and 36.1% ± 18.0% respectively) (Figure 1B). Following these experiments, 1.5 mg/kg scopolamine and 10 mg/kg VU846 were chosen as optimal doses for the larger study investigating the effect of an M1-PAM on LM in M1-PD mice.

**Figure 1.**
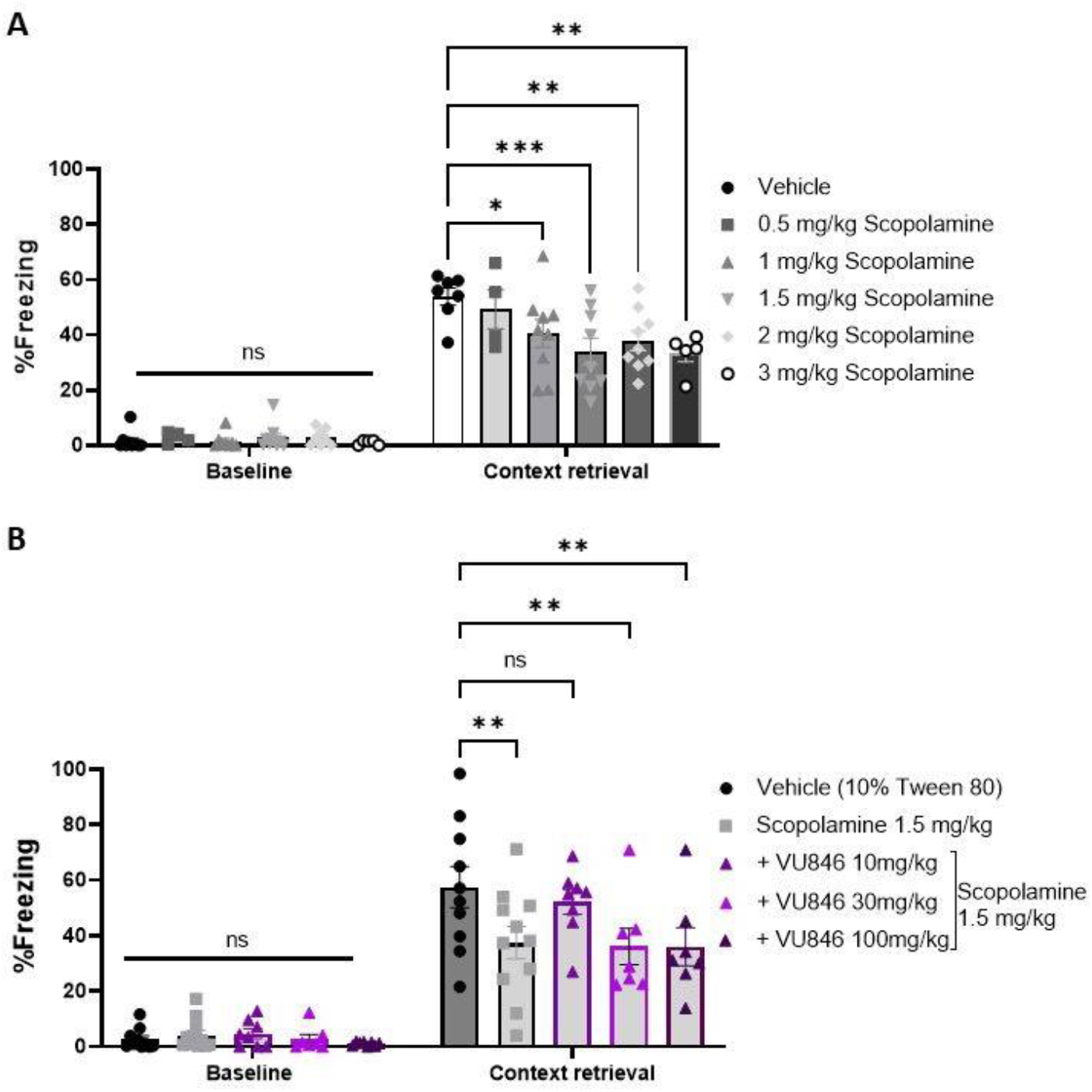
Optimisation of scopolamine and M1-PAM doses in M1-WT mice. M1-WT mice were treated with varied doses of scopolamine (n=4-9) **(A)**, or subsequently 1.5 mg/kg scopolamine with varied doses of the M1-PAM VU846 (n=7-11) **(B**), 30 mins prior to training in a fear conditioning paradigm and then returned to the same environment 24 hrs later for contextual memory retrieval whereby a reduced level of freezing indicates impaired memory. 2-way ANOVA with Dunnett’s post-hoc correction for multiple comparisons compared to vehicle, *p<0.05, **p<0.01, ***p<0.001, ns=not significant.

### Contextual memory retrieval in phospho-deficient M1 mice

In contextual memory retrieval, scopolamine induced a deficit in M1-WT, M1-KO and M1-PD mice reducing the freezing response in response to the context from 71.9% ± 12.3%, 63.3% ± 17.8% and 30.2% ± 16.1% to 41.0% ± 18.4%, 34.2% ± 18.1% and 16.2% ± 8.4% respectively (Figure 2A-C). This effect, however, was partially rescued by the M1-PAM VU846 only in M1-WT mice (freezing level with VU846 treatment 56.0% ± 14.6%) (Figure 2A) and not when M1 was absent (Figure 2B) or lacking phosphorylation sites (Figure 2C). When results were compared between strains, looking at the vehicle treated groups only, M1-PD mice displayed a deficit in their freezing response to contextual memory retrieval compared to both M1-WT and M1-KO mice suggesting that biased M1 signalling causes impairment of memory, while a lack of M1 is insufficient to cause this deficit alone (Figure 2D). This result was further corroborated using novel object recognition (NOR) testing where M1-PD mice displayed a cognitive deficit compared to M1-KO mice (Supplemental Figure 1). The M1-PD receptor has previously been shown to have impaired internalisation (Bradley et al., 2020; Scarpa et al., 2021) and approximately 30% increased maximal receptor activity (Bradley et al., 2020) compared to M1-WT. Further to this, the M1-PD receptor shows a differential staining pattern in hippocampal tissue with evidence of perinuclear staining of M1 receptor in M1-WT which is absent in M1-PD animals (Figure 2E). Furthermore, this internalisation is constitutive and occurs in the absence of receptor activation in mice expressing M1-DREADD receptor (Designer Receptor Exclusively Activated by Designer Drugs) where the staining pattern was similar to M1-WT (Figure 2E). This suggests that internal cellular localisation of the M1 receptor is dependent on phosphorylation and points to localisation of the receptor at the membrane as the cause of resulting phenotype.

**Figure 2.**
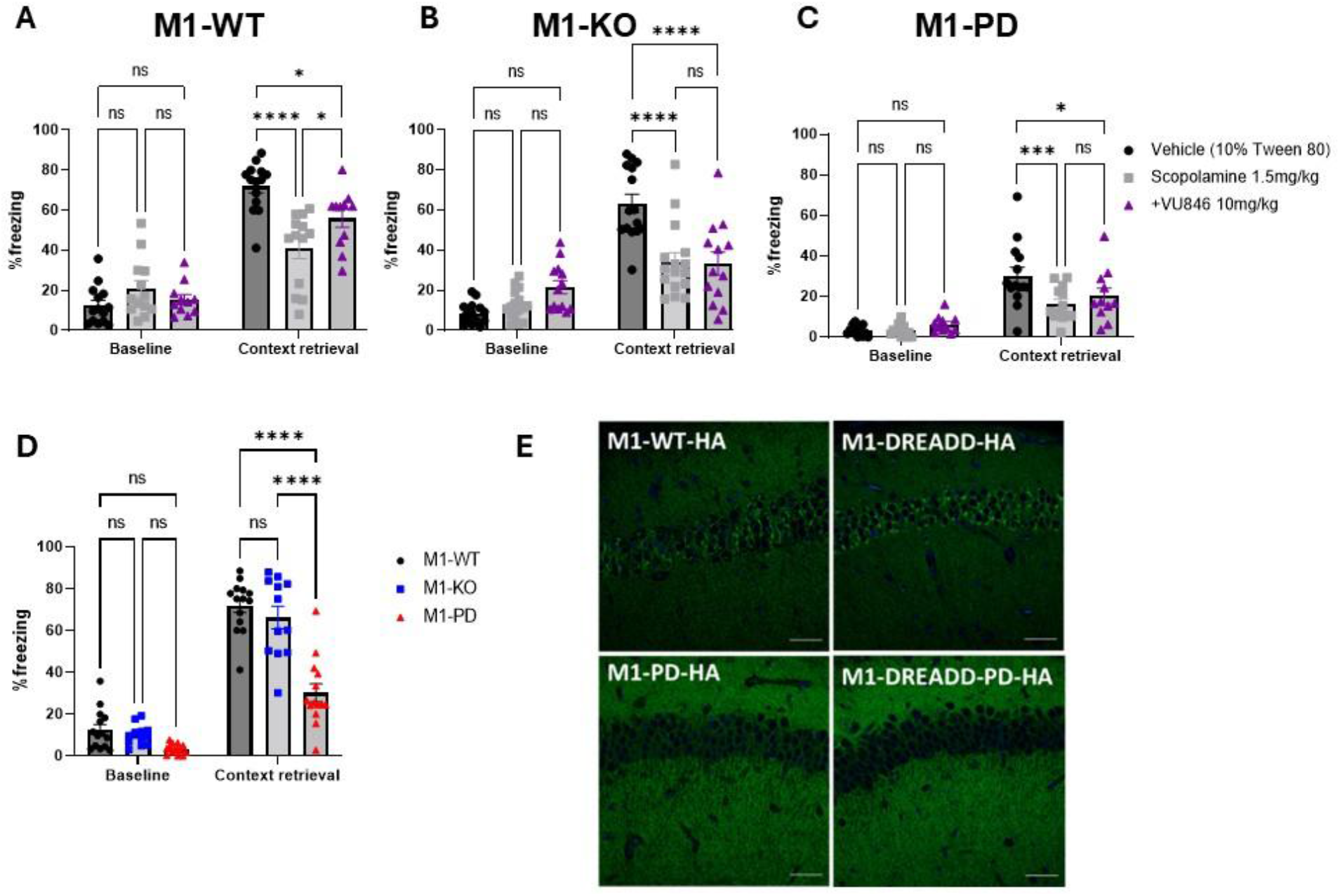
M1-PAM rescues scopolamine induced learning and memory deficit in contextual memory retrieval in M1-WT but not M1-KO or M1-PD mice, and untreated M1-PD but not M1-KO mice display a deficit compared to WT. M1-WT-HA (n=11-14) **(A)**, M1-KO (n=13-16) **(B)** or M1-PD-HA (n=11-14) **(C)** mice were treated with 1.5 mg/kg scopolamine +/-10 mg/kg VU846 30 mins prior to training in a fear conditioning paradigm. Mice were then returned to the same environment 24 hrs later for contextual memory retrieval whereby a reduced level of freezing indicates impaired memory. The response in vehicle treated animals across strains was also compared **(D)**. 2-way ANOVA with Tukey’s post-hoc correction for multiple comparisons, *p<0.05, ***p<0.001, ****p<0.0001, ns=not significant. **(E**) Representative images of CA1 hippocampal region of HA-tagged M1-WT, M1-DREADD, M1-PD and M1-DREADD-PD stained with HA antibody demonstrating location of the receptor (green). Images at 40X magnification, scale bar = 50 µm

### Cued memory retrieval in phospho-deficient M1 mice

In cued memory retrieval, scopolamine again induced a deficit in M1-WT mice, reducing the freezing response to the tone from 81.9% ± 13.5% to 61.5% ± 20.3%, and this was rescued by VU836 treatment where the freezing response increased to 75.6%± 14.9% (Figure 3A). In M1-KO and M1-PD mice, however, scopolamine did not induce any deficit in freezing response (Figure 3B,C). This suggests that the M1 receptor is crucially important in cued memory retrieval as the effect of scopolamine on the other muscarinic receptors does not impair the LM response. Furthermore, corroborating this, when vehicle treated animals were compared across strains both M1-KO and M1-PD mice displayed a deficit in their response compared to M1-WT, with a freezing response of 65.6% ± 14.5% and 42.8% ± 20.1% respectively (M1-WT 81.9% ± 13.5%) (Figure 3D). M1-PD mice, however did also display a reduction in their baseline freezing response to the tone compared to M1-WT which may suggest a non-memory related phenotypic difference resulting in this particular observation (baseline values M1-WT 31.5 % ±24.9%, M1-PD 10.2% ±8.2%) (Figure 3D).

**Figure 3.**
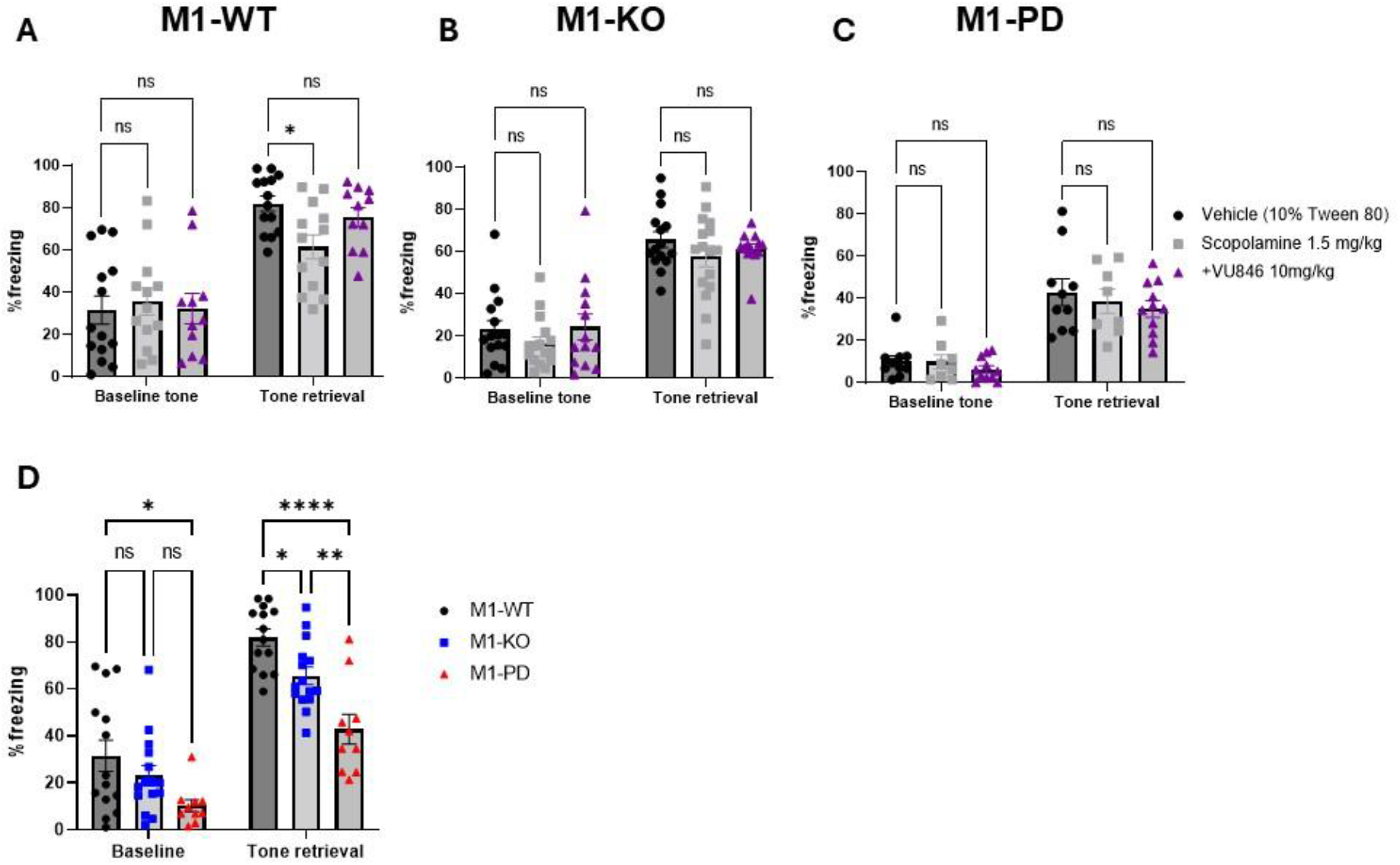
M1-PAM rescues scopolamine induced learning and memory deficit in cued memory retrieval in M1-WT but scopolamine does not induce a deficit in M1-KO or M1-PD mice, and both untreated M1-PD and M1-KO mice display a deficit compared to WT. M1-WT-HA (n=11-14) **(A)**, M1-KO (n=13-16) **(B)** or M1-PD-HA (n=8-11) **(C)** mice were treated with 1.5 mg/kg scopolamine +/-10 mg/kg VU846 30 mins prior to training in a fear conditioning paradigm. Mice were then returned to a new environment the following day and presented with the tone for cued memory retrieval. 2-way ANOVA with Dunnett’s post-hoc correction for multiple comparisons to vehicle treatment, *p<0.05, ns=not significant. **(D)** The response in vehicle treated animals across strains was also compared. 2-way ANOVA with Tukey’s post-hoc correction for multiple comparisons, *p<0.05, ***p<0.001, ****p<0.0001, ns=not significant.

Together, these data therefore suggest that lack of phosphorylation of the M1 receptor results in impaired recognition and associative memory, while lack of M1 receptor specifically impairs only cued memory retrieval, and that phosphorylation of the M1 receptor is required for any LM benefits associated with M1 agonism.

### Orthosteric vs allosteric M1 agonism in fear conditioning memory

Although the biased M1-PD receptor impaired memory, enhancement of endogenous ACh M1 signalling using an M1-PAM did not cause further impairment in the M1-PD mice (Figure 4A). However, in an additional mouse line where the M1 receptor was globally replaced by a chemogenetic M1-DREADD receptor (Designer Receptor Exclusively Activated by Designer Drugs), activation of this receptor prior to fear conditioning with clozapine-n-oxide (CNO) significantly and robustly impaired contextual memory retrieval compared to vehicle treated mice (Figure 4B), with a similar effect also observed in cued retrieval (Figure 4C). Meanwhile WT mice showed no effect with CNO suggesting this is a direct result of M1 agonism and not clozapine off target effects (Figure 4D).

**Figure 4.**
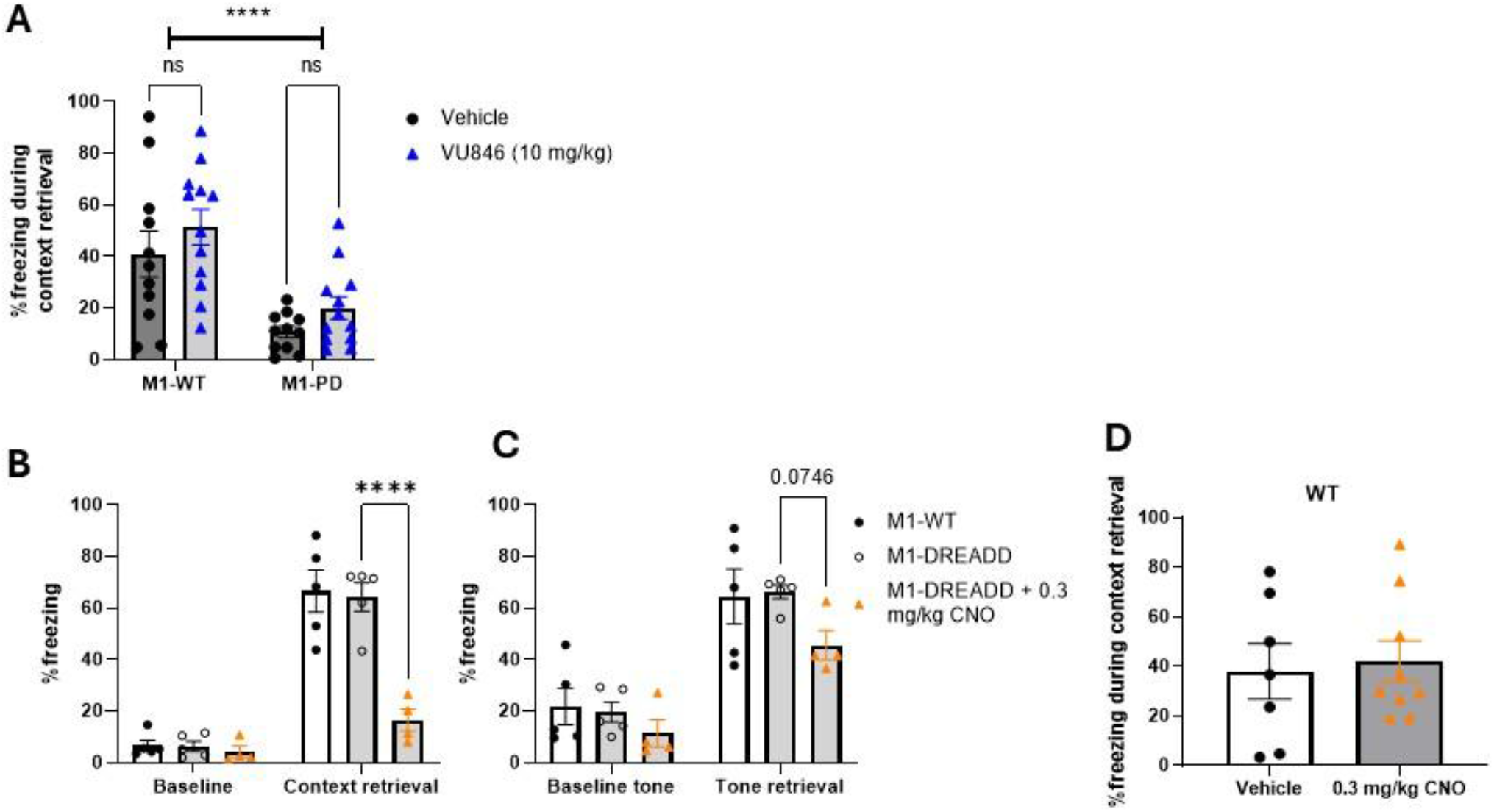
Orthosteric agonism of M1-DREADD receptor impairs memory but allosteric agonism of M1-WT or M1-PD receptor does not. **(A)** Male and female M1-WT-HA (n=11) and M1-PD (n=12) mice were treated with the M1-PAM VU846 30 mins prior to fear conditioning and returned to the chamber 24 hrs later for contextual memory retrieval. 2-way ANOVA ****p<0.0001 effect of strain, no significant effect of drug overall p=0.12, Bonferroni’s post-hoc correction for multiple comparisons between vehicle and VU846 for each strain, ns= not significant. **(B & C)** Male M1-DREADD mice were treated with 0.3 mg/kg of clozapine-N-oxide (CNO), an orthosteric agonist of the DREADD receptor, or vehicle (n=4-5 per group) 30 mins prior to training in a fear conditioning paradigm and then returned to the same environment 24 hrs later for contextual memory retrieval **(B)** or presentation of the tone in an altered environment 48 hrs later for cued memory retrieval **(C)** whereby a reduced level of freezing indicates impaired memory. **(D)** WT mice (n=7-9 per group) were also treated with 0.3 mg/kg CNO in a separate experiment demonstrating no effect of CNO. 2-way ANOVA with Dunnett’s post-hoc correction for multiple comparisons, ****p<0.0001, all other comparisons p>0.05.

Taken together these results suggest that *abhorrent* M1 signalling, whether from direct orthosteric drug activation or lack of internalisation of the receptor, may impair memory while enhancement of endogenous M1 signalling does not.

## Discussion

The results presented here demonstrate an importance of phosphorylation of the M1 receptor for LM, and highlight the importance of finding the balance of an optimal dose when it comes to targeting the M1 receptor pharmacologically for cognition or LM, as abhorrent M1 signalling appears to impair memory.

Scopolamine induced a deficit in LM in M1-WT mice similar to that seen in C57Bl/6J WT mice previously with 1.5 mg/kg inducing an approximately 50% reduction in context dependent freezing (Bradley et al., 2018) suggesting no impact of the HA-tag on the receptor’s behaviour in this task. This is not surprising considering an M1 receptor with a larger monomeric enhanced green fluorescent protein (mEGFP) tag also displayed similar receptor activity to the WT receptor (Marsango et al., 2022). This deficit was rescued by 10 mg/kg of the M1-PAM, VU846, but not by higher doses, which is not dissimilar to previous results with another M1-PAM, BQCA, whereby scopolamine rescue by the PAM demonstrated a bell-shaped curved response (Ma et al., 2009). Indeed, other investigations of scopolamine LM rescue by M1 orthosteric ligands have also displayed this bell-shaped curve with the optimal dose often not being the highest dose (Bradley et al., 2018). Importantly, this result was reproducible in the main study whereby VU846 rescued the scopolamine-induced deficit in both contextual and cued fear memory retrieval in M1-WT mice with no rescue apparent in context retrieval for either M1-KO or M1-PD mice. The former confirms the specificity of VU846 at the M1 receptor, and the latter confirms the need for phosphorylation of the receptor for this effect. The deficit induced by scopolamine across the three strains was comparable with a 43%, 46% and 47% reduction in context dependent freezing for the M1-WT, M1-KO and M1-PD mice respectively, which does suggest some redundancy regarding the M1 receptor in this particular LM assessment as this pan-muscarinic antagonist could still induce a robust deficit in the absence of M1 receptor. This is further supported by the lack of a deficit in contextual memory retrieval in the M1-KO mice compared to M1-WT suggesting M1 is not essential. Indeed, numerous studies have implicated the G_q/11_ coupled M3 muscarnic receptor in fear memory response (da Silva et al., 2022; Diniz et al., 2022; Fedoce et al., 2016; Poulin et al., 2010). Nevertheless, the data does confirm that, even if contextual memory retrieval is not entirely dependent on M1 signalling, a deficit in this regard can be rescued with an M1-specific PAM and for this to be effective, phosphorylation of the M1 receptor is required.

Scopolamine did not induce a deficit in cued fear memory retrieval in either M1-KO or M1-PD mice but only in M1-WT mice and furthermore, M1-KO and M1-PD mice displayed a deficit in cued memory retrieval when vehicle treatment groups were directly compared to M1-WT mice. This indicates that M1 is required for cued memory retrieval, which is in line with previous reports, utilising a very similar fear conditioning testing paradigm, where a deficit was demonstrated in M1-KO mice in cued but not contextual memory retrieval (Miyakawa et al., 2001). However, although M1 is apparently only specifically required for cued memory retrieval, phosphorylation of M1, and therefore appropriate M1 signalling, has implications for various forms of memory indicated here by deficits in M1-PD mice in both cued and contextual fear memory and the NOR test. Contextual fear memory and NOR are known to be primarily dependent on the hippocampus (Cohen et al., 2013; Phillips & LeDoux, 1992; Wallenstein & Vago, 2001), while cued fear conditioning memory also involves the amygdala (Phillips & LeDoux, 1992). Early evidence indicates that while muscarinic receptors M1, M3 and M4 are detectable in the hippocampus of the rat, only M1 is detectable in the amygdala (Buckley et al., 1988). This may explain the differences observed, where the amygdala and therefore cued memory retrieval is severely impacted by M1-KO. Hippocampal memory, however, appears to only be affected when M1 signalling is abhorrent through the lack of internalisation of the receptor in the hippocampus (Figure 2E) but not impacted by receptor KO due to contributions of compensatory signalling. This theory is further supported by observations that M3-KO mice display deficits in context retrieval but not cued retrieval (Poulin et al., 2010), which suggests that a lack of M1 expression could be compensated by M3 in context retrieval but not cued retrieval.

Differences in locomotor activity due to M1-KO could of course account for difference observed in freezing behaviour as previously suggested (Miyakawa et al., 2001). However, distance travelled was not significantly greater in M1-KO mice compared to M1-WT mice during any stage measured (Supplemental figure S2A) which suggests that this deficit observed in cued retrieval is indeed due to a memory deficit. Previously, M1-PD mice have demonstrated a hypolocomotive response in open field and elevated plus maze testing environments but not when locomotion was measured throughout the 24hr cycle using telemetry (Bradley et al., 2020). Interestingly, the mice in the current study displayed hyperlocomotion compared to WT and M1-KO mice in the fear conditioning chamber environment (Supplemental figure S2A). Chronically stressed mice can display hyperlocomotion in the presence of acute stressors (Strekalova et al., 2005) and therefore it is possible that the fear conditioning chambers caused additional stress to the already anxious mice (Bradley et al., 2020) resulting in this observed hyperlocomotion and potentially the reduced freezing response. However, an analysis of strain variations in fear conditioning response suggested little correlation between locomotor activity and freezing level (Seemiller et al., 2021) and linear regression analysis revealed no significant correlation of freezing level and locomotor activity of the M1-PD mice at baseline (Supplemental figure S2B). The NOR test, however, is less sensitive to locomotor phenotypes because a minimum investigation time threshold can be used to exclude animals which do not explore objects due to hyper- or hypolocomotion (Lueptow, 2017). Therefore, considering object investigation times were similar across all strains (Supplemental Figure 1B), we can be quite confident that the deficit observed in M1-PD mice in this test suggests a memory deficit.

Taken together, and despite the potential locomotion limitations, these data do suggest that lack of phosphorylation of the M1 receptor results in an impairment in reference memory overall, while lack of the M1 receptor impacts cued fear memory only. Previous work investigating the signalling of M1-PD showed little difference to the WT receptor except for a decrease in β-arrestin coupling, reduced internalisation of the receptor and an increase in maximum response (Bradley et al., 2020; Scarpa et al., 2021). Considering the largest effect appears to be on receptor internalisation, this suggests this may be the root cause of phenotypic differences. Indeed, a distinctly different pattern of staining of the M1 receptor was observed between both M1-WT and M1-PD, and M1-DREADD and M1-DREADD-PD mice, with perinuclear staining absent in both phosphodeficient strains, confirming that receptor localisation differs between phosphorylation-intact and phosphorylation-deficient receptors. The ability to internalise the receptor may perhaps lead to specified localisation of the receptor which is necessary for appropriate LM signalling, for example numerous GPCRs have been shown to signal from internalised locations such as endosomes or the Golgi apparatus (Plouffe et al., 2020). In addition, without internalisation to “switch off” the receptor signalling, and levels of receptor expression being equivalent between strains (Bradley et al., 2020; Scarpa et al., 2021), this may lead to an excess of M1 signalling as demonstrated by increased levels of G protein activation and second messenger (IP_3_) accumulation ex vivo in M1-PD mice (Bradley et al., 2020). As previously discussed, a bell-shaped curve is often observed in studies investigating M1 receptor agonism in LM (Bradley et al., 2018; Brown et al., 2021; Ma et al., 2009; Rook et al., 2018). This may support the hypothesis that a balance in M1 signalling needs to be achieved for optimal LM effects, with excessive M1 signalling, due to lack of receptor internalisation, being detrimental to LM. This is further supported by an observed LM deficit caused by activation of the M1-DREADD receptor during the fear conditioning testing paradigm at 0.3 mg/kg but not at lower doses (Supplemental figure S3). This is a direct orthosteric activation of the receptor and suggests that direct M1 activation results in a profound effect on LM. Considering the M1-DREADD mice display no deficit in LM compared to WT, this direct M1 activation would be in combination with whatever compensatory signalling mechanisms are taking place and could perhaps be over-activation. In contrast, however, boosting the endogenous ACh signalling in M1-PD mice with VU846 did not impair LM further, which may suggest that the timing of M1 signalling is also critical and that it is excess or abhorrent signalling which results in this detriment.

The data presented here suggests that phosphorylation of the M1 receptor is crucially important for LM potentially due to the requirement of internalisation of the receptor either to switch off the signal and avoid excessive signalling or the need for specific cellular localisation of the receptor for appropriate signalling. This adds further support to previous work suggesting the need to design M1 ligands which are not G_q_ biased, to ensure phosphorylation of the receptor occurs for appropriate LM signalling (Bradley et al., 2020). In addition, it highlights the need to consider the dose of M1 ligands carefully when used in studies investigating LM in both preclinical and clinical subjects which is of key importance considering the interest in this receptor for treating dementia

## Methods

### Animals

Knock-in mouse lines expressing Ha-tagged M1-WT (n=66), HA-tagged phosphodeficient M1 receptor (M1-PD) (n=54), Ha-tagged M1-DREADD receptor (n=14) (Bradley et al., 2020) or lacking M1 receptor (M1-KO) (n=63) were bred as homozygotes onto a C57BL/6J background. Mice were fed ad libitum with a standard mouse chow and were maintained in the appropriate animal facility at least one week prior to experimentation on a standard 12 hr light/dark cycle. Mice were used for behavioural experiments between 8-18 weeks of age and acclimatised to the behavioural testing room overnight prior to testing and to the experimenter in the week prior to testing. All experiments were carried out in the morning by the same experimenter. All studies were carried out in accordance with the Animals Scientific Procedures Act 1986 and under an appropriate project licence held by Andrew Tobin.

### Drugs and dosing

The drugs utilised in this study were the pan-muscarinic antagonist, to induce a learning and memory (LM) deficit, scopolamine hydrobromide (Sigma) at 0.5-3 mg/kg doses; the M1 positive allosteric modulator (PAM) VU0486846 (VU846) at 10-100 mg/kg; and the M1-DREADD activating ligand clozapine-n-oxide (CNO) (Tocris) at 0.3 mg/kg. Mice were randomised to receive drug or vehicle (10% Tween-80 or 5% glucose) by intraperitoneal injection by one experimenter 30 mins prior to undergoing a fear conditioning paradigm by a different blinded experimenter.

### Fear conditioning learning and memory test

Male mice were placed in a conditioning chamber (Stoelting ANY-maze Fear Conditioning System) for a period of 6.5 mins. This fear conditioning paradigm consisted of a baseline 2 min period followed by 3 successive tone-foot shock pairings spaced 1 min apart whereby the tone (conditioned stimulus, 2.8 kH, 85 dB) played for 30 sec and the foot shock (unconditioned stimulus, 0.4 mA) coincided with the final 2 sec of tone. After the final tone-shock pairing the mice remained in the chamber for 1 min. The following day, mice were returned to the same conditioning chamber for a period of 3 min and the percentage time spent freezing was measured using AnyMaze to assess context dependent memory retrieval. The following day from this, mice were returned to the conditioning chambers where the chambers were modified with opaque patterned walls and an orange scent to create an alternative environment. Mice remained in these modified chambers for 2 min before being presented with the same conditioned stimulus as day 1 (tone) for 2 mins and the percentage time spent freezing was measured using AnyMaze to assess cued dependent memory retrieval.

### Novel object recognition learning and memory test

Female mice were habituated to 60 cm diameter opaque buckets for periods of 5 mins, 10 mins, 15 mins and 18 mins over 4 successive days. On the 5^th^ day mice underwent novel object recognition (NOR) in the same buckets whereby mice were allowed to explore two identical objects, either 2X towers of Duplo® blocks or 2X 50 mL Duran pyrex bottle half filled with corncob bedding for a period of 18 mins (exploration phase). After 2.5 hrs, mice were returned to the same bucket where the mouse was presented with both a familiar object (previously seen bottle or block) and novel object (previously unseen bottle or block) and allowed to explore for 10 mins (testing phase). The time spent investigating each object was measured during both phases using AnyMaze software whereby an investigation zone of 2.5 cm was drawn around the object. A discrimination index was then calculated where a positive value indicates more time spent investigating the novel object:

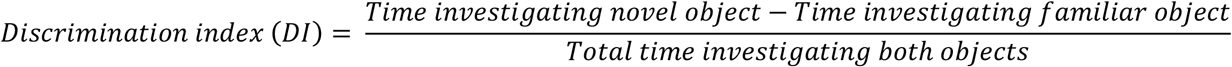

### Immunohistochemistry

Mice expressing HA-tagged M1-WT, M1-PD, M1-DREADD or M1-DREADD-PD were culled by transcardiac perfusion of phosphate buffered saline followed by 4% paraformaldehyde (PFA). Brains were removed and post-fixed in 4% PFA for at least 24 hrs before embedding in 2.5% agarose and sectioning into 50 μm coronal sections using a vibratome (Leica). Sections containing the hippocampus were stained using rabbit anti-HA (1/500, Cell signalling, C29F4) overnight at 4 °C followed by a secondary antibody (1/500, anti-rabbit Alexa Fluor 488) and imaged using confocal microscopy at 40X magnification.

### Statistical analysis and exclusions

All behavioural testing was performed by an experimenter blinded to the drug group and genotype but analyses were performed using AnyMaze to further minimise bias. Sample size calculations for the fear conditioning and NOR were performed using preliminary data (Minitab) and suggested, to detect differences of 24% freezing and a DI of 0.2 respectively with 80% power, that group sizes of n=12 and n=11 were required respectively. All data was analysed using GraphPad Prism 9.2 using the statistical tests outlined in the figure legends. A p value of <0.05 was considered significant. Some mice were excluded from the analysis for the following reasons: n=7 (spread over M1-KO, M1-WT-HA, M1-DREADD) due to technical failure of the fear conditioning foot shock system; n=4 (M1-KO) due to technical failure during NOR; n=8 (spread over M1-WT-HA, M1-KO, M1-PD-HA) due to being identified as having a high level of baseline freezing when first exposed to the fear conditioning chamber (ROUT method of outlier identification Q=1%); n=1 (M1-WT-HA) due to observed stereotypical behaviour on the day of testing.

## Supplemental figures

**Supplemental figure 1.**
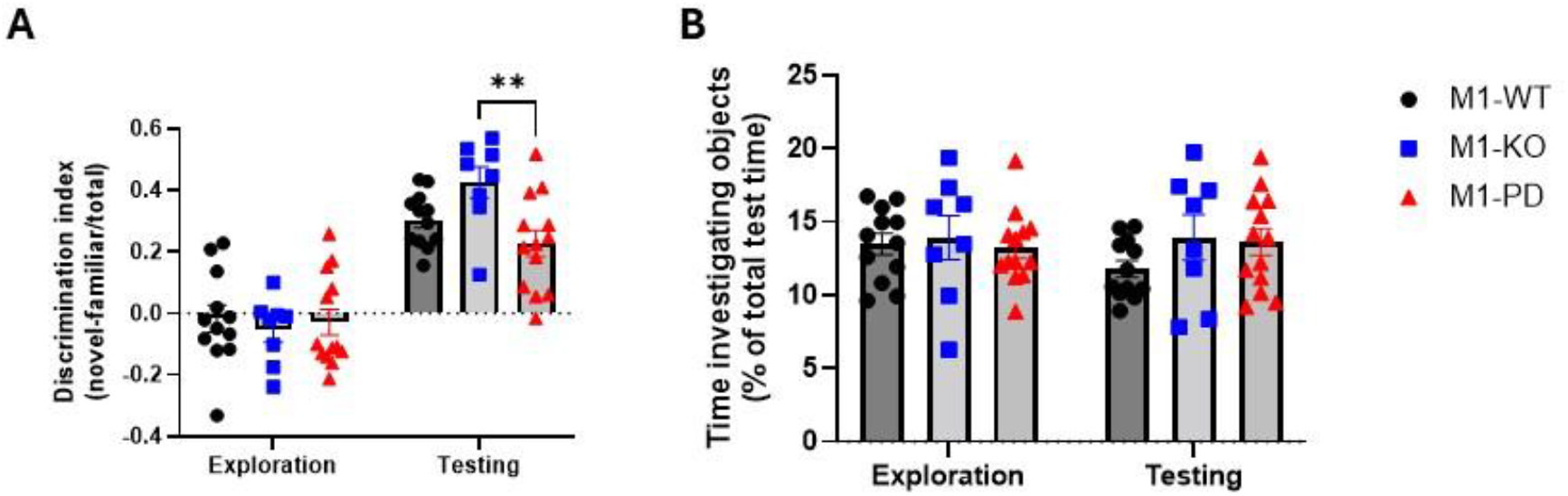
M1-PD mice display a deficit in novel object recognition. Female M1-WT, M1-KO and M1-PD mice (n=8-13) were assessed in novel object recognition and a discrimination index showing proportion of time spent investigating the novel object in the testing phase, indicating intact memory, was calculated **(A)**. The proportion of time spent investigating both objects during both exploration and testing was also measured to validate the result **(B)**. 2-way ANOVA with Tukey’s post-hoc correction for multiple comparisons, **p<0.01. All other comparisons not significant.

**Supplemental figure 2.**
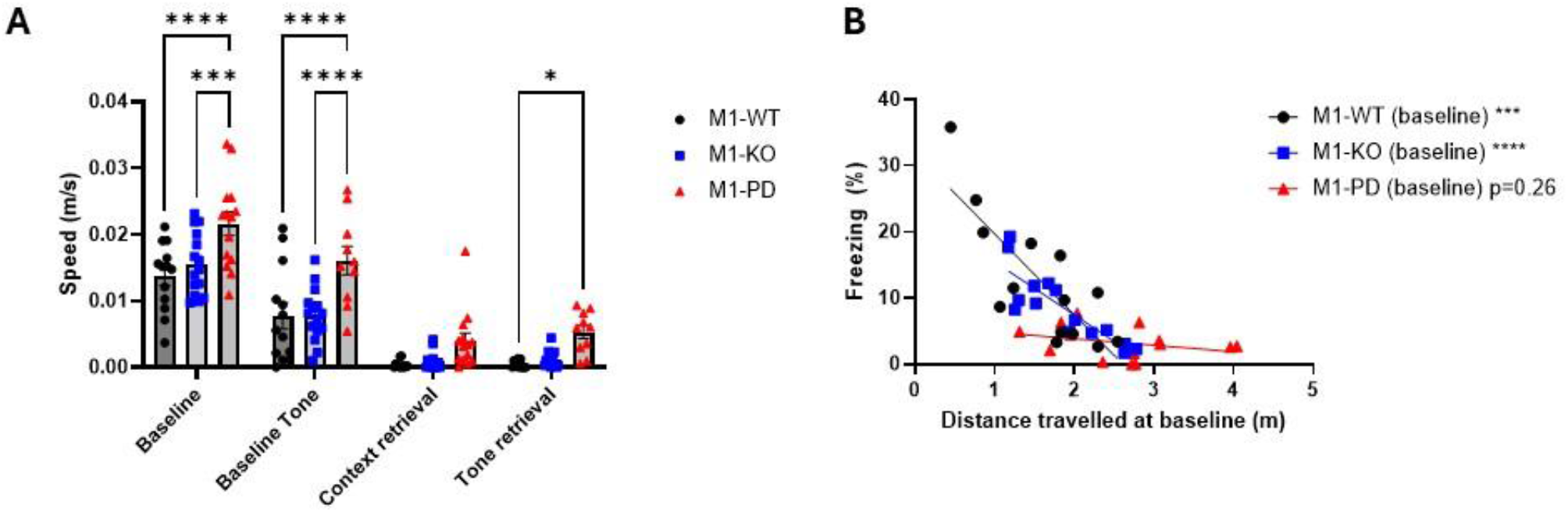
Locomotion measures of vehicle treated M1-WT, M1-KO and M1-PD mice during fear conditioning testing paradigm. **(A)** Distance travelled by mice during training periods (baseline & baseline tone) and retrieval periods (context & cued) presented as speed to account for differences in the duration of each period. 2-way ANOVA with Tukey’s post-hoc correction for multiple comparisons, *p<0.05, ***p<0.001, ****p<0.0001. **(B)** Linear regression analysis of distance travelled during baseline period and freezing level during baseline period where stars and p value indicate the deviation of the slope from zero. This suggests that there is no correlation between freezing level and locomotion in M1-PD mice.

**Supplemental figure 3.**
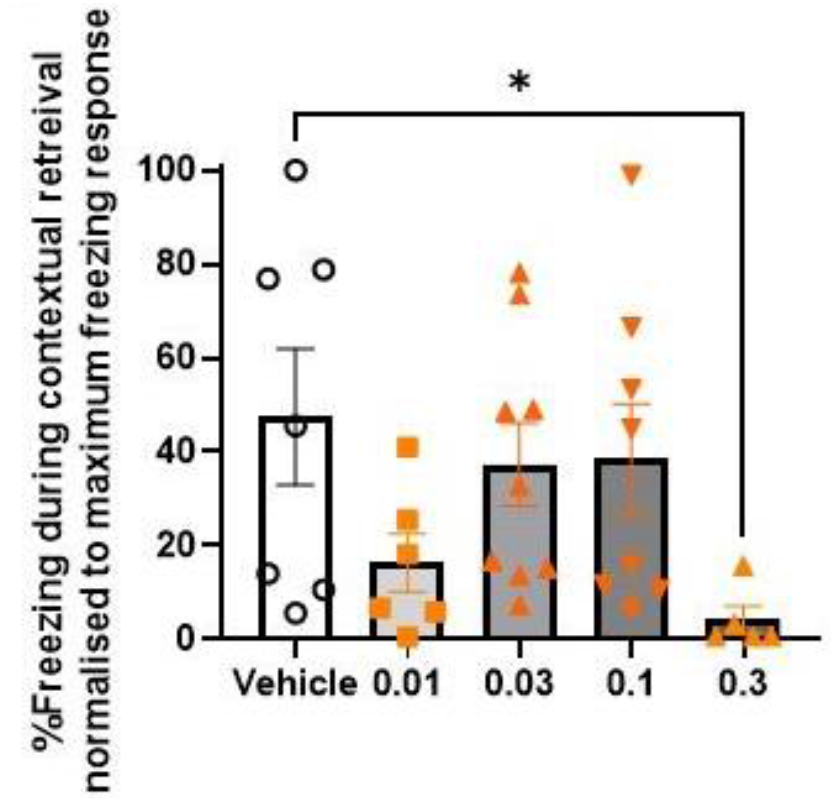
Contextual memory retrieval in M1-DREADD mice with increasing doses of CNO. Male M1-DREADD mice were treated with 0.01 - 0.3 mg/kg of clozapine-N-oxide (CNO), an orthosteric agonist of the DREADD receptor, or vehicle (n=5-9 per group) 30 mins prior to training in a fear conditioning paradigm and then returned to the same environment 24 hrs later for contextual memory retrieval. Data presented as freezing level normalised to the highest freezing response. One way ANOVA with Bonferroni’s post hoc correction for multiple comparisons to vehicle treatment group *p<0.05. All other comparisons not significant.

